# A human impact metric for coastal ecosystems with application to seagrass beds in Atlantic Canada

**DOI:** 10.1101/457382

**Authors:** Grace E.P. Murphy, Melisa Wong, Heike K. Lotze

**Affiliations:** Department of Biology, Dalhousie University, 1355 Oxford Street, P.O. Box 15000, Halifax, Nova Scotia, Canada, B3H 4R2; Fisheries and Oceans Canada, Bedford Institute of Oceanography, 1 Challenger Drive, Dartmouth, Nova Scotia, Canada, B2Y 4A2

**Keywords:** anthropogenic threats, human impact, coastal ecosystems, biogenic habitat, seagrass, *Zostera marina*, coastal management, conservation planning

## Abstract

Coastal biogenic habitats are particularly vulnerable to cumulative human impacts from both terrestrial and marine realms. Yet the broad spatial scale used in current global or regional approaches of quantifying multiple anthropogenic stressors are not relevant to the local or bay-wide scales affecting most coastal biogenic habitats. To fill this gap, we developed a standardized human impact metric to quantify the magnitude of anthropogenic impacts to coastal ecosystems more broadly, and biogenic habitats in particular. We applied this metric to 180 seagrass beds (*Zostera marina*), an important biogenic habitat prioritized for marine protection, across Atlantic Canada. Our impact metric includes five bay-scale and four local-scale terrestrial and marine-based impacts. Results show that seagrass beds and coastal bays in Atlantic Canada exist across a wide gradient of human impacts. Considerable differences in the range and intensity of impacts within and between regions provide insight into where coastal bays and seagrass ecosystems are expected to be most and least affected by individual or cumulative human threats. We discuss implications for management and conservation planning, and the general application of our impact metric to other coastal regions and habitats in Canada and beyond.

## Introduction

Over past decades and centuries, the magnitude, spatial extent and variety of human impacts have substantially increased in coastal ecosystems around the world (Lotze *et al.* 2006, Halpern *et al.* 2008). Nearshore biogenic habitats, such as seagrass meadows, kelp forests, rockweed beds, and oyster reefs, are especially vulnerable as they are subject to anthropogenic threats from both the terrestrial and marine realms (Orth *et al.* 2006, Worm and Lotze 2006, Waycott *et al.* 2009, Beck *et al.* 2011, Krumhansl *et al.* 2016). Coastal management strategies have recognized nearshore biogenic habitats as areas of high conservation value, and their inclusion in marine protected areas (MPAs) is a conservation priority worldwide (DFO 2007, Cullen-Unsworth and Unsworth 2016). Despite this, it remains unclear how to prioritize areas for protection given the multitude of anthropogenic stressors impacting these ecosystems. Metrics of anthropogenic stressors used to inform management and conservation have previously been applied to ocean ecosystems across broad global and regional scales (Halpern *et al.* 2008, Ban and Alder 2008, Murray *et al.* 2015). However, these assessments are not relevant at smaller spatial scales, such as specific coastal bays, nearshore ecosystems or biogenic habitats.

Impact metrics useful for management and conservation planning of coastal ecosystems, particularly for specific biogenic habitats, should quantify impacts at both bay-wide and local scales. Human impacts are widely recognized as having scale-dependent effects on ecosystem processes (Powell *et al.* 2013), and spatial scale may also be relevant when assessing the magnitude of stressors influencing marine ecosystems (Thrush *et al.* 1999). For example, human activities were found to influence seagrass ecosystems mainly at the local scale, typically within 1-3km of the seagrass bed (Shelton *et al.* 2017, Cullain *et al.* 2018a, Iacerella *et al.* 2018). Yet, given that coastal biogenic habitats are often confined to bays and estuaries they are also influenced by stressors that operate at the bay-scale, such as nutrient and sediment loading and pollution run-off from the surrounding watersheds (McIver *et al.* in review). Additionally, various biogenic habitat types may be differentially affected by human activities, suggesting the need for habitat-specific assessments. A comprehensive assessment of human impacts in coastal ecosystems should thus consider impacts to specific habitats at both the local and bay-wide scale.

In Canada, eelgrass (*Zostera marina*) has been designated an Ecologically Significant Species (ESS), and the inclusion of eelgrass beds within MPA networks is a central conservation priority (DFO 2009a). Given their global distribution and critical role in ecosystem functioning (Duarte 2002, Hughes *et al*. 2009), eelgrass beds are an ideal case study for the development of a human impact metric relevant for coastal ecosystems. Global declines in seagrass cover and associated ecosystem health has been attributed to various human activities, including nutrient pollution (Orth *et al.* 2006), spread of invasive species (Wong and Vercaemer 2012, Williams 2007), coastal land alteration (Grech *et al.* 2012), construction of overwater structures (Fresh *et al.* 2006, Thom *et al.* 2011), and aquaculture (Skinner *et al.* 2013, Cullain *et al.* 2018a). While natural events are also sometimes responsible for large-scale seagrass losses (i.e., ice scour, storms; Duarte 2002), human activities are recognized as a significant driver of seagrass ecosystem degradation (Short and Wyllie-Echeverria 1996, Hemminga and Duarte 2000, Waycott *et al.* 2009). A metric that quantifies multiple human impacts at spatial scales relevant for seagrass beds will aid conservation planning by identifying priority areas with low human impacts and highlight areas where management measures should be considered.

The objectives of this study were to (1) develop a general standardized human impact metric that can quantify the magnitude and range of anthropogenic impacts on various coastal ecosystems and biogenic habitats, and (2) apply this metric to seagrass beds in Atlantic Canada. First, we selected relevant anthropogenic impacts known to influence biogenic habitats, in particular seagrass beds, on local or bay-wide scales based on the published literature. We then compiled empirical data for these impacts for 180 seagrass beds inhabiting 52 bays across two biogeographic regions in Atlantic Canada and assessed the distribution of these impacts across the different seagrass beds, coastal bays, and biogeographic regions. Finally, we explored the potential general utility of our human impact metric for other biogenic habitats, coastal ecosystems, and geographic locations to inform coastal management and conservation.

## Methods

### General human impact metric development

We began by selecting relevant anthropogenic impacts for inclusion in the human impact metric for coastal ecosystems and biogenic habitats. Here, we focused on impacts known to influence seagrass beds on local or bay-wide scales and relevant indicators or measurable proxies based on the published literature (Table 1). We then assessed data availability to quantify the extent of each impact across our study region. To standardize the human impact metric across all sites, we focused on impacts and indicators for which comparable data were available from the three relevant provincial governments (Nova Scotia, New Brunswick, Prince Edward Island) or the two marine management regions of Fisheries and Oceans Canada (Maritimes region and Gulf region), which are also considered separate biogeographic regions (Scotian Shelf and Gulf of St. Lawrence, respectively) with unique oceanographic and hydrodynamic processes (Spalding *et al.* 2007, DFO 2009b).

**Table 1.**
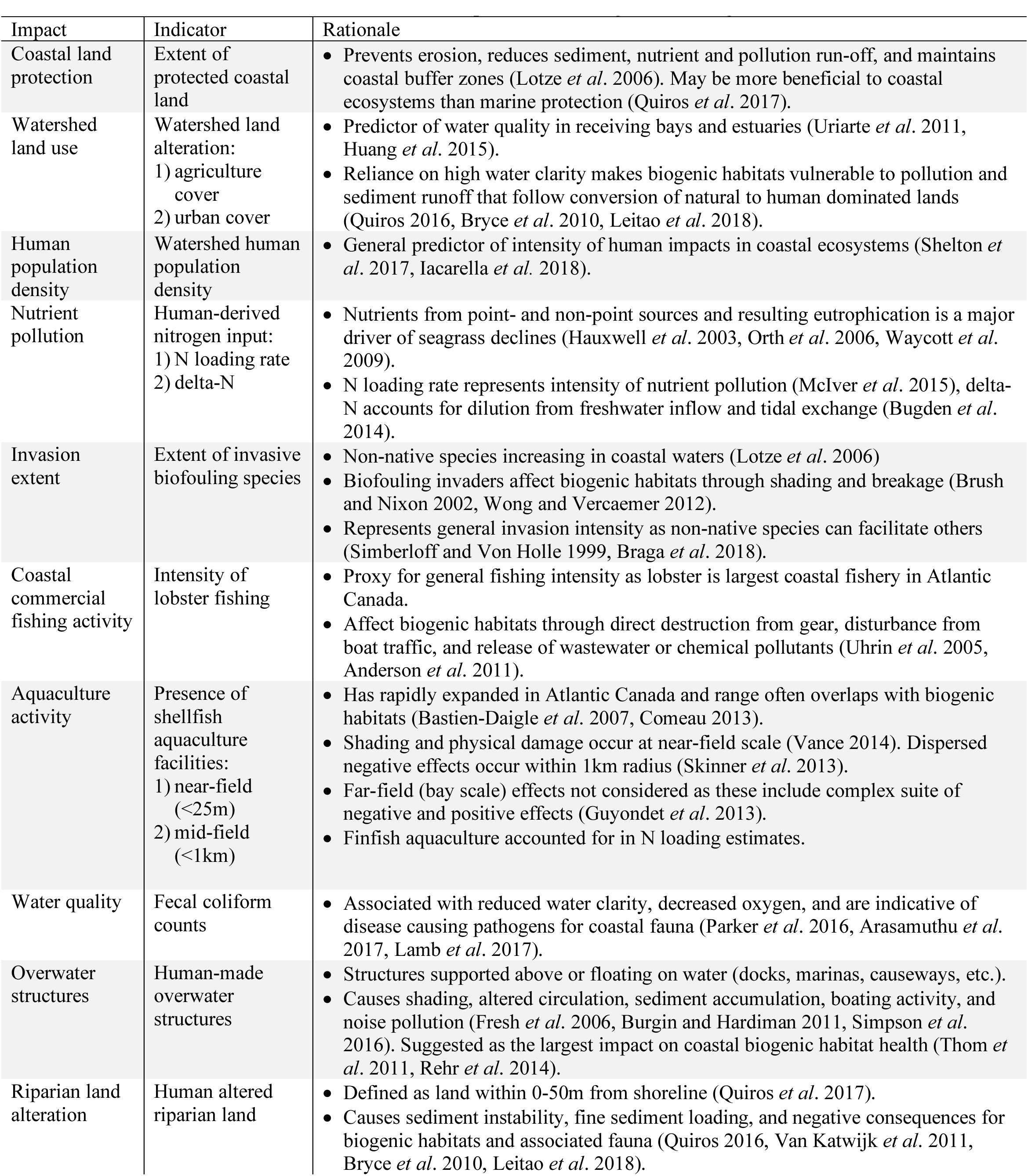
Rationale for selection of human impacts influencing coastal biogenic habitats

We separated impacts into those acting at a bay-scale or local-scale (0-1km). The bay-scale was defined by the geographic boundary of the bay (or estuary) and considered impacts that influence the entire waterbody of the bay, for example through tidal mixing or indirect effects, thereby also influencing the biogenic habitats within. In our case, this included watershed land use, human population density, nutrient pollution, invasion extent, and coastal commercial fishing activity (Fig. 1). In contrast, local-scale impacts were defined as those more directly affecting the biogenic habitat, whereby the considered local range depends on both the impact and the habitat in question. For our seagrass beds, we included aquaculture activity, water quality, overwater structures, and riparian land alteration (Fig. 1) within 0-1km of the habitat. This smaller scale has been suggested most relevant for assessing human impacts to seagrass beds (Shelton *et al.* 2017, Iacarella *et al.* 2018).

**Figure 1.**
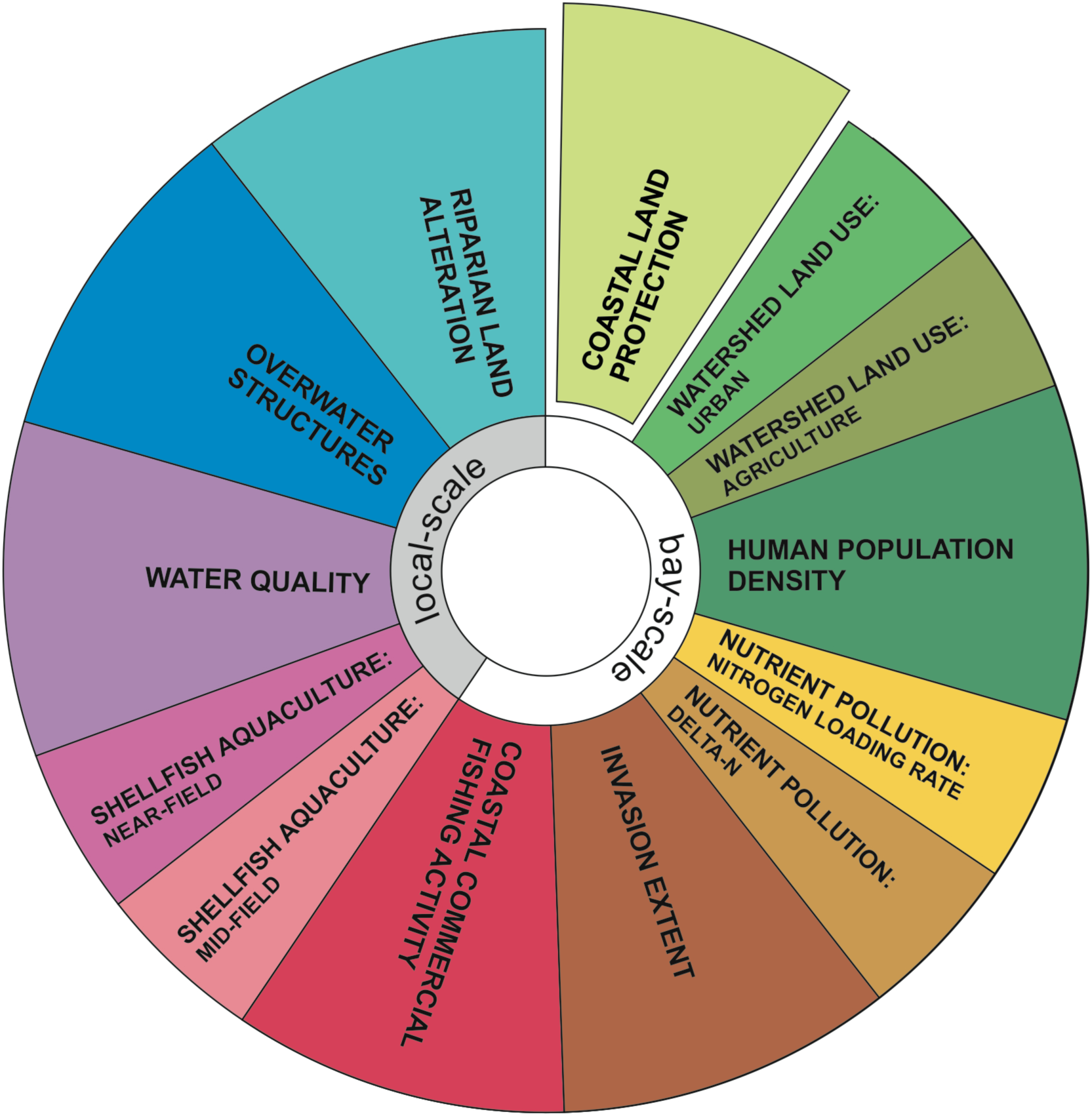
All bay- and local-scale human impacts included in the standard human impact metric for seagrass beds in Atlantic Canada. Note that coastal land protection represents a benefit as opposed to a stressor. Also note that three impacts (land use, nutrient pollution, and aquaculture) were further separated into sub-measures.

Our derived human impact metric therefore included five bay-scale and four local-scale impacts, with three of these impacts (land use, nutrient pollution, and aquaculture activity) each further subdivided, resulting in a total of 12 impact scores (Fig. 1). In addition, we included coastal land protection as a measure expected to benefit our biogenic habitat. Depending on the coastal ecosystem or biogenic habitat in question, these impacts and indicators and the scale at which they are assessed can be easily adapted. See Table 1 for detailed rationale for the inclusion of these impacts in our human impact metric for seagrass beds in Atlantic Canada.

### Data collection

Our goal was to apply the human impact metric to seagrass beds across Atlantic Canada. To do so, we first compiled the locations of 180 seagrass beds (Fig. 2) from field surveys conducted over the past decade (Schmidt *et al.* 2012, Skinner *et al.* 2013, Cullain *et al.* 2018b, Wong 2018, Weldon *et al.* 2009, Locke and Bernier (unpublished data)) in 52 bays along the coasts of Nova Scotia (NS), New Brunswick (NB), and Prince Edward Island (PEI) and in two biogeographic regions (Scotian Shelf, Gulf of St. Lawrence).

**Figure 2.**
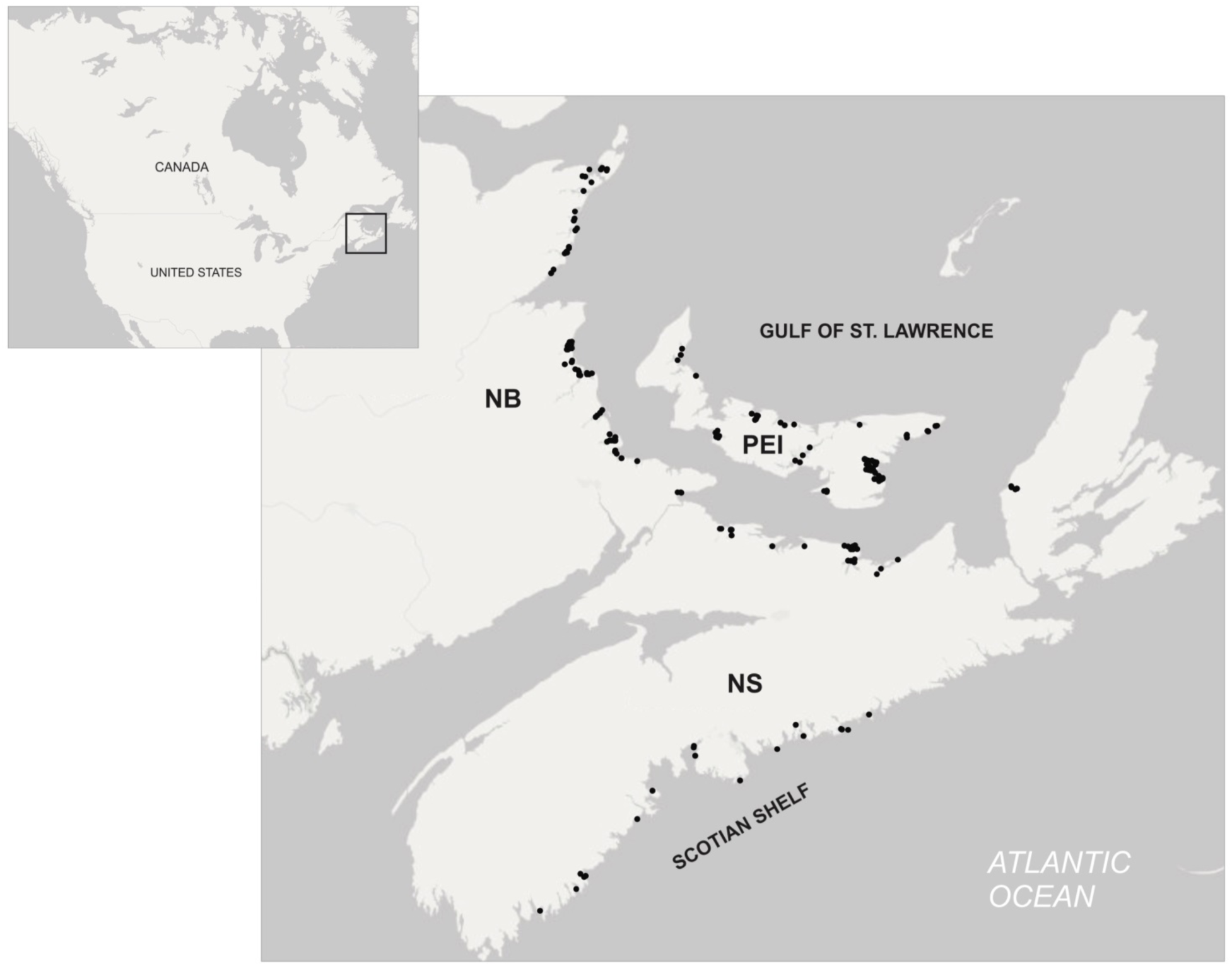
Location of 180 seagrass sites (black dots) that the human impact metric was applied to in three Atlantic Canadian provinces (NS = Nova Scotia, NB = New Brunswick, PEI = Prince Edward Island) and two biogeographic regions (Scotian Shelf, Gulf of St. Lawrence).

#### Coastal land protection

We obtained GIS shapefiles of provincial, federal, and private conservation areas from Environment and Climate Change Canada. We clipped shapefiles to the 0-200m coastal land surrounding each bay to determine the percentage of protected land within each bay’s coastal land zone. We defined coastal land as 0-200m from the coastline because this distance has previously been identified as important for pollution input to receiving waters (Valiela *et al.* 1997).

#### Land use

We used previously delineated watershed boundaries for each bay provided by provincial authorities (NS Department of Environment, NB Department of Energy and Resource Development, PEI Department of Land and Communities). In some cases, we further delineated watersheds to include all freshwater inputs by predicting bay-specific watercourse drainage patterns using hydrographic data and digital terrain models (GeoNova, GeoNB, PEI GIS Data Catolog). We clipped shapefiles that classified land use types across each of the provinces to the predetermined watershed areas for each bay. The percentage urban and agriculture land use were determined by summing appropriate units in the watershed areas and dividing by total watershed area. We also determined the number of bays with >10% combined urban and agriculture watershed land use as this threshold has been identified as presenting a moderate to high risk for ecosystem degradation in receiving waters (WWF 2017) and has been used to differentiate developed vs. undeveloped watershed land (Lerberg et al. 2000, Bilkovic et al. 2006, Blake *et al.* 2014).

#### Human population density

Watershed human population density was estimated using the number of civic addresses present in each watershed (GeoNova, GeoNB, PEI GIS Data Catolog) multiplied by the average number of residents per household (2.3, 2.3, and 2.4 in NS, NB, and PEI, respectively) (Statistics Canada 2017a). The number of individuals was standardized to watershed area.

#### Nutrient pollution

Nutrient pollution was divided into two sub-measures: nitrogen loading rate and delta-N (ΔN). We used data from previous applications of a nitrogen loading model (NLM), originally developed for Waquoit Bay, Massachusetts (Valiela *et al.* 1997), for several bays in Atlantic Canada, including 7 bays in NB (McIver *et al.* 2015), 21 bays in NS (Nagel *et al.* 2018), and 6 bays in PEI (Palmer 2018). For the remaining 19 bays, we estimated total nitrogen load (kg N yr^−1^) using linear regression models specific to each province (Supplementary Methods, Fig. S1 and S2). Briefly, the NLM estimates watershed and atmospheric derived total dissolved nitrogen loads from several point (i.e., direct atmospheric deposition, wastewater discharge, seafood processing plants, finfish aquaculture) and non-point sources (i.e., indirect atmospheric deposition, septic systems, and fertilizer addition) with appropriate loss parameters (Valiela *et al.* 1997, McIver *et al.* 2015, McIver *et al.* 2018). Final nitrogen loading rates were calculated by summing across all sources and standardized to the area of each bay (kg N ha bay^−1^ yr^−1^) to account for dilution effects. We assessed how many bays had nitrogen loading rates above the 50 kg N ha bay^−1^ yr^−1^ threshold identified by Latimer and Rego (2010) as being detrimental to seagrass coverage.

The ΔN combines total nitrogen load (kg N yr^−1^, as calculated above) with a bay’s tidal flushing time and freshwater recharge volume to estimate the increase in nitrogen concentration above ambient oceanic nitrogen concentration after dilution factors (Bugden *et al.* 2014). We calculated tidal flushing and freshwater recharge for each bay as described in Nagel *et al.* (2018). We assessed how many bays had ΔN values above the 0.06 threshold identified by Bugden *et al.* (2014) as being likely to experience anoxic events as a result of excess nitrogen input.

#### Invasion extent

We used presence/absence data from the Aquatic Invasive Species (AIS) monitoring program (Sephton *et al.* 2017) to estimate invasion extent for each bay. This program monitors the presence of nine invasive biofouling species by assessing settlement on plastic monitoring plates. These species included seven tunicates: vase tunicate (*Ciona intestinalis*), clubbed tunicate (*Styela clava*), carpet tunicate (*Didemnun vexillum*), golden star tunicate (*Botryllus schlosseri*), violet tunicate (*Botrylloides violaceous*), compound sea squirt (*Diplosoma listerianum*), and European sea squirt (*Ascidella aspersa*), a bryozoan (*Membranipora membranacea*) and an amphipod, the Japanese skeleton shrimp (*Caprella mutica).* While the European green crab (*Carcinus maenas*) is also invasive in our study region and can negatively impact seagrass beds (Garbary *et al.* 2014), we were unable to obtain quantitative estimates of green crab densities for each of our sites to include in our quantification of invasion extent but note that green crabs are known to occur in all 52 bays.

Our metric of invasion extent is a measure of invasive biofouling species richness over 10 years. We used a 10-year average (2006-2015) of invader presence/absence for each of the nine invasive biofouling species and averaged across species to obtain a measure of invasion extent for each bay. When AIS monitoring stations were not present in a bay we used the closest station within 50km. When no monitoring station was located within 50km for a specific year we excluded that year from the 10-year average. Since the AIS monitoring stations are not located within seagrass beds, our measure of invasion extent is a proxy for how likely fouling is expected within a bay and is not indicative of invaders specifically fouling seagrass at that site.

#### Coastal commercial fishing activity

We used data from Fisheries and Oceans Canada to calculate the number of inshore and offshore lobster fishing licences per port in 2018 as an estimate of lobster fishing intensity (A. Cook, pers. comm. 2018). While these data do not indicate where lobster fishing is taking place (i.e. inshore, offshore, or in an adjacent bay), they provide a general estimate of the extent of fishing vessel traffic.

#### Aquaculture activity

Shellfish aquaculture activity was measured as presence/absence. We used data from the NS Department of Fisheries and Aquaculture, the NB Department of Agriculture, Aquaculture, and Fisheries, and Fisheries and Oceans Canada to determine the presence or absence of operational shellfish aquaculture leases in 2018 within <25m radius (near-field) and <1km radius (mid-field) of each seagrass bed. The impact of finfish aquaculture was included in nutrient loading estimates as outlined above.

#### Water quality

We used fecal coliform monitoring data from the Canadian Shellfish Sanitation Program (CSSP 2016) as a measure of water quality. We used a 10-year average (2005-2015) of fecal coliform counts (Most Probable Number [MPN] 100ml^−1^] from the CSSP monitoring station closest to each seagrass bed, which were typically <500m away. We assessed water quality contamination according to thresholds set by CCSP as fecal coliform counts <14 MPN 100ml^−1^ considered uncontaminated and of good quality (CSSP 2016).

#### Overwater structures

We manually classified the total area of overwater structures (i.e., wharfs, bridges, causeways, etc.) within a 1km radius of each seagrass bed using Google Earth (Google Inc. 2018). We standardized the total area of overwater structures to the water surface area within a 1km radius.

#### Riparian land alteration

Using the land use data described above, we estimated human altered riparian land within 0-50m from the coastline due to urban, agriculture, and forestry land use in contrast to unaltered natural wetland, forest or conservation land areas. We define riparian land as 0-50m from the coastline because this distance has previously been used to define riparian land and human land alteration within this range has been shown to be detrimental to seagrass health (Quiros *et al.* 2017). We measured the area of riparian land alteration within a 1km radius of seagrass, which was standardized to the total land area within the 1km radius of each seagrass site. We used the same land alteration thresholds for watershed land use (>10%) to assess how many seagrass beds had adjacent riparian land considered a moderate to high risk for ecosystem degradation in receiving waters (WWF 2017). This threshold is based on total watershed land alteration, as opposed to strictly riparian land, as no threshold estimates exist for riparian land alteration. Given that coastal land alteration near seagrass beds is expected to be more detrimental than alteration in the entire watershed (Quiros *et al.* 2017), we expect this to be a conservative threshold.

### Human impact standardization

We used the above data to first calculate the intensity of each human impact at each of the 180 seagrass beds (raw data). Each impact was then standardized to range from 0-1, and the multiple impacts at each site were compiled into a petal diagram to illustrate the overall human impact level. Standardization was performed at two different spatial scales: across all 52 bays or 180 sites and across each region.

## Results

Comparing the intensity of human impacts across all 52 coastal bays (Fig. 3) and 180 seagrass beds (Fig. 4) revealed a wide gradient of human impacts in Atlantic Canada. We observed differences in the distribution of each impact between the Scotian Shelf and Gulf of St. Lawrence, and within the Gulf region between the mainland coast (NB and NS) and PEI (Fig. 3 and 4 insets). Below we describe and assess these differences among regions (Atlantic NS, Gulf NB+NS, and PEI) based on 95% confidence intervals (Fig. 3 and 4 insets).

**Figure 3.**
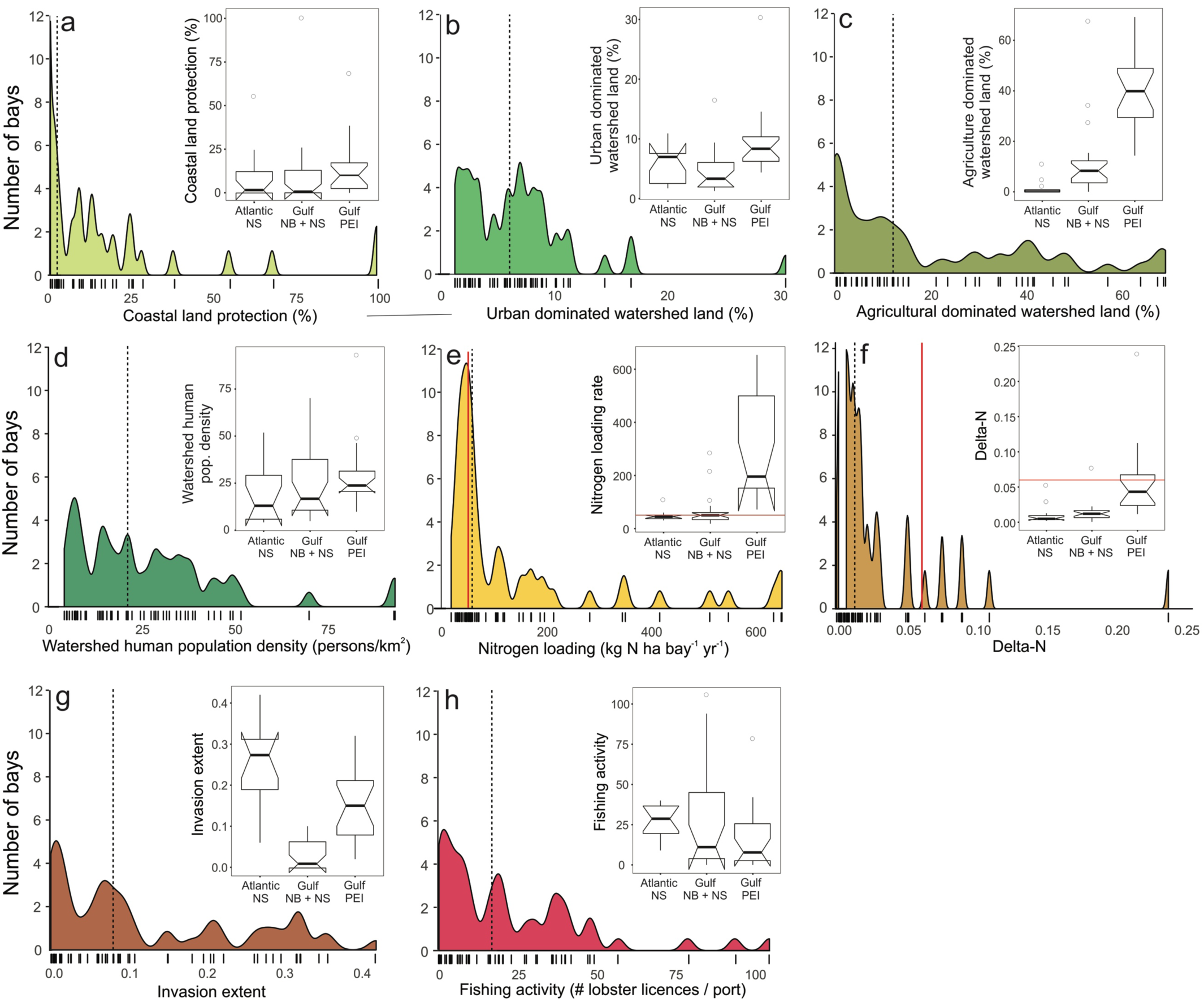
Distributions and notched boxplots of coastal land protection and bay-scale human impact scores for all bays (n=52). Scores are raw values as opposed to the regionally-standardized scores displayed in Figures 5 and 6. Dashed vertical lines indicate the median score for each impact. Red lines in (e) nitrogen loading and (f) delta-N distributions and box plots indicate threshold levels identified in the literature. Tick marks on the x-axes indicate the scores of individual observations. Boxplots show the median, the first and third quartiles, and outliers, and notches indicate 95% confidence intervals for medians. Note different scales on axes.

**Figure 4.**
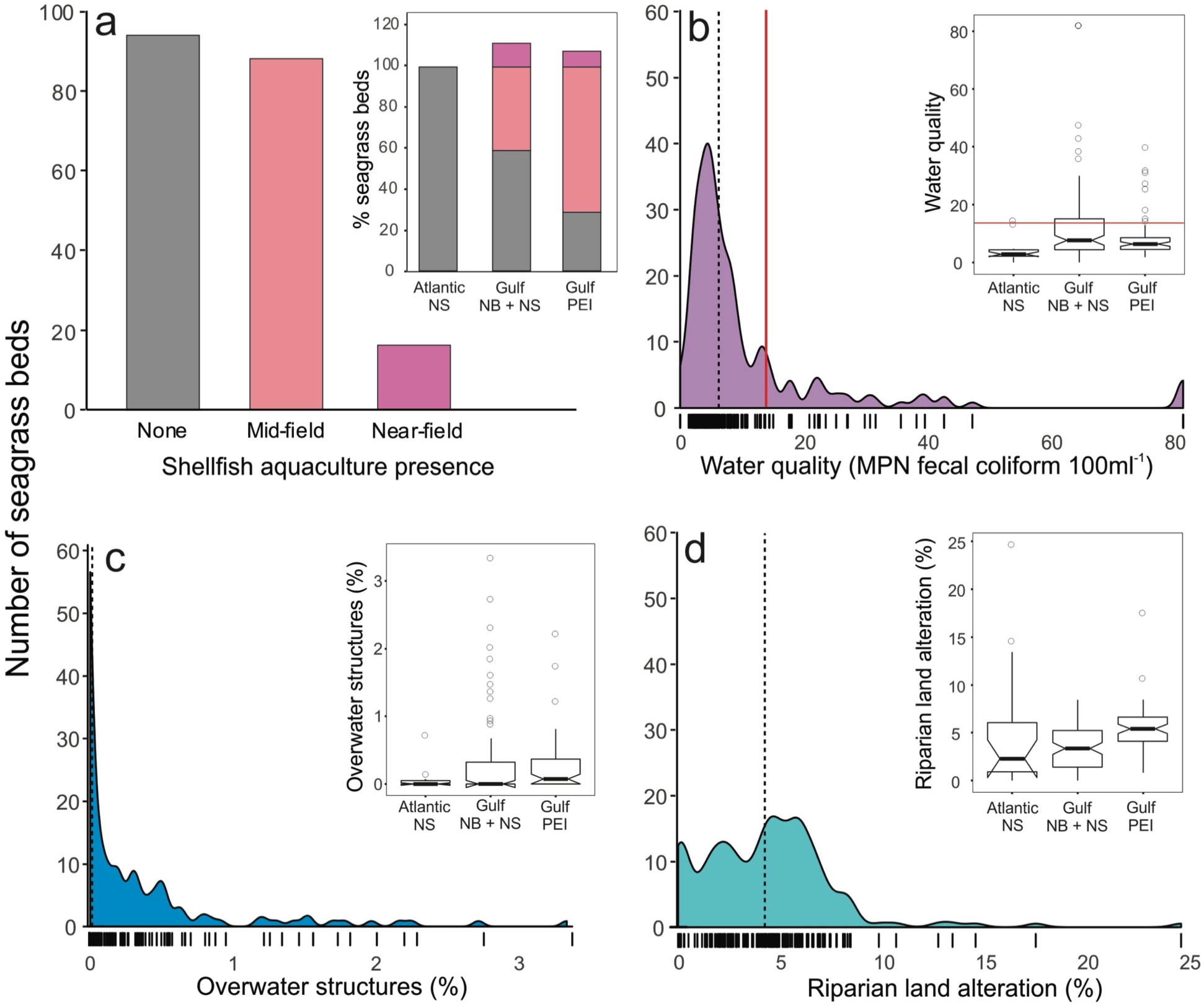
Distributions and notched boxplots of local-scale human impact scores for all seagrass beds (n=180). Scores are raw values as opposed to the regionally-standardized scores displayed in Figures 5 and 6. Dashed vertical lines indicate the median score for each impact. Red lines in (b) water quality degradation distribution and box plot indicate contamination threshold level based on CSSP. Tick marks below each distribution indicate the scores of the individual observations. Boxplots show the median, the first and third quartiles, and outliers. Notches in boxplots show 95% confidence intervals for medians. Note that in (a) inset the percentage can be greater than 100% as seagrass beds with both near-field and mid-field aquaculture facilities are included. Note different scales on axes.

### Bay-scale impacts

#### Land protection and use

The percentage of coastal land protection was highly variable, ranging from 0% to 100% protection (median = 3.21%; Fig. 3a). Coastal land protection was highest in PEI (10.04% ± 5.42% 95%CI), much greater than in Gulf NB+NS (0.68 ± 4.32%) and Atlantic NS (1.65 ± 5.53%), although not statistically significant. PEI watersheds also had the greatest land alteration for urban (Fig. 3b) and agricultural uses (Fig. 3c), although urban land use was similar between PEI and Atlantic NS (8.3 ± 1.49% and 7.1 ± 2.23%, respectively), but significantly greater than Gulf NB+NS (3.5 ± 1.35%). Agricultural land use in PEI (39.8 ± 7.11%) was significantly greater than in the Gulf NB+NS (8.4 ± 2.87%) and Atlantic NS (3.3 ± 0.32%). According to the threshold of >10% watershed land alteration (WWF 2017), 65% of the 52 bays were at risk for ecosystem degradation in receiving waters, most of them in Gulf NB+NS and PEI, and only 2 in Atlantic NS.

#### Human population density

Similar to the regional variation in land protection and use, the median human population density was higher in PEI (24.1 persons km^−2^) than in Atlantic NS (13.2 km^−2^) and Gulf NB+NS (16.9 km^−2^), although not statistically different among regions (Fig. 3d).

#### Nutrient pollution

The median human derived nitrogen loading rate across the 52 bays was 58.9 ± 25.9 kg N ha bay^−1^ yr^−1^. Nitrogen loading rates were significantly higher in PEI bays (193.7 ± 122.22) than in Atlantic NS (45 ± 5.84) and Gulf NB+NS (52.5 ± 8.83; Fig. 3e). Across all bays, 64% were above the 50 kg N ha bay^−1^ yr^−1^ threshold identified by Latimer and Rego (2010) as being detrimental to seagrass coverage (Fig. 3e). This included all (100%) of bays in PEI, 55% in Gulf NB+NS, but only 25% in Atlantic NS. A similar pattern was observed for ΔN, indicating that PEI bays had the highest eutrophication risk (0.041 ± 0.016; Fig. 3f), followed by Gulf NB+NS bays (0.012 ± 0.003), although not significantly different, while ΔN in Atlantic NS bays (0.005 ± 0.003) was significantly lower than in both PEI and Gulf NB+NS. Only 15% of all bays had ΔN values above the 0.06 threshold that would indicate high probability of anoxic events (Bugden *et al.* 2014). Again, most bays above this threshold were located in PEI while only one Gulf NB+NS bay (River Philip) and no Atlantic NS bays were considered at risk for anoxic events.

#### Invasion extent

The invasion extent of non-native biofouling species was significantly higher in Atlantic NS bays (median = 0.28 ± 0.05; Fig. 3g). Invasion extent in PEI (0.15 ± 0.05) was also significantly higher than in Gulf NB+NS (0.01 ± 0.02). The dominant invaders across the three regions were vase tunicates, violet tunicates, golden star tunicates, and *Membranacia membranipora.* However, *M. membranipora* was not detected in any PEI bays, and European sea squirts were only detected in Atlantic NS bays.

#### Coastal commercial fishing activity

Lobster fishing was significantly higher in Atlantic NS bays (median = 29 ± 7.6 licenses port^−1^) than in PEI (8 ± 8.4), while that in the Gulf NB+NS (11 ± 13.5) was not significantly different from the other regions (Fig 3h).

### Local-scale impacts

#### Aquaculture activity

Near-field shellfish aquaculture (<25m from seagrass bed) was present at 20 seagrass beds and mid-field (<1km) at 87 beds across all 180 sites (Fig. 4a). Among regions, Atlantic NS had no shellfish aquaculture within 1km of any seagrass beds, and mid-field aquaculture was more prevalent in PEI (71% of beds) and Gulf NB+NS (41%) than near-field (8% and 15% respectively).

#### Water quality

Across all 180 sites, fecal coliform counts close to seagrass beds ranged from 0-82 MPN 100ml^−1^ (median = 6.1 ± 1 MPN 100ml^−1^, Fig 4b). Median fecal coliform counts near seagrass beds were significantly lower in Atlantic NS (2.58 ± 0.8 MPN 100ml^−1^) than in PEI (6.1 ± 0.8) and Gulf NB+NS (7.62 ±1.7; Fig. 4b), indicating that, overall, water quality is uncontaminated and of good quality according to the CCSP threshold (<14 MPN 100ml^−1^). However, 35 (19%) of seagrass beds had water quality above this threshold, with 1 in Atlantic NS, 9 in PEI and 25 in Gulf NB+NS.

#### Overwater structures

The percentage of water covered by overwater structures near seagrass beds ranged from 0-3.38% across all seagrass beds (Fig 4c). The median percentage was greater in PEI (0.074 ± 0.07%) than in Gulf NB+NS (0.00 ± 0.03%) and Atlantic NS (0.00 ± 0.02%), with no statistical differences among regions (Fig. 4c).

#### Riparian land alteration

Similar to human land alteration in the entire watershed, the median percentage of coastal land altered in close proximity to seagrass beds was significantly higher in PEI (5.4 ± 0.5%, Fig. 4d) than in Atlantic NS (2.3 ± 1.9%) and Gulf NB+NS (3.3 ± 0.6%). However, the range of coastal land alteration was greater in Atlantic NS (0-25%) compared to PEI (1-18%) and Gulf NB+NS (0-8%). Of the 180 seagrass beds, 3% were affected by >10% riparian land alteration. Most of these were in Atlantic NS, accounting for nearly 25% of all Atlantic NS seagrass beds.

### Human impacts at different spatial scales

Our human impact metric can be compared across all coastal bays and seagrass beds in Atlantic Canada (Fig. 3 and 4) as well as within biogeographic regions (e.g. Scotian Shelf and Gulf of St. Lawrence, Fig. 5 and 6). Standardizing the impacts within each region allows for better comparison of the importance of each impact relative to other sites subjected to similar biogeographic conditions. For example, compared to all other seagrass beds, those on the Atlantic coast of NS had relatively low human impact (Fig. 3 and 4 insets); yet when standardized within the Scotian Shelf region, relative impact patterns became more apparent (Fig. 5). This revealed that several seagrass beds were minimally impacted by human activities relative to other Atlantic NS sites, including Port Joli, Cable Island and Taylor’s Head, which have higher coastal protection and lower overall bay and local-scale impacts (Fig. 5). In comparison, other seagrass beds were more impacted by human activities, with Second Peninsula highly impacted by urban and agriculture land use, poor water quality, and commercial fishing, St. Margaret’s Bay by riparian land alteration, poor water quality, and commercial fishing, Sambro by high population density and urban land use, and Musquodoboit by nutrient pollution (Fig. 5).

**Figure 5.**
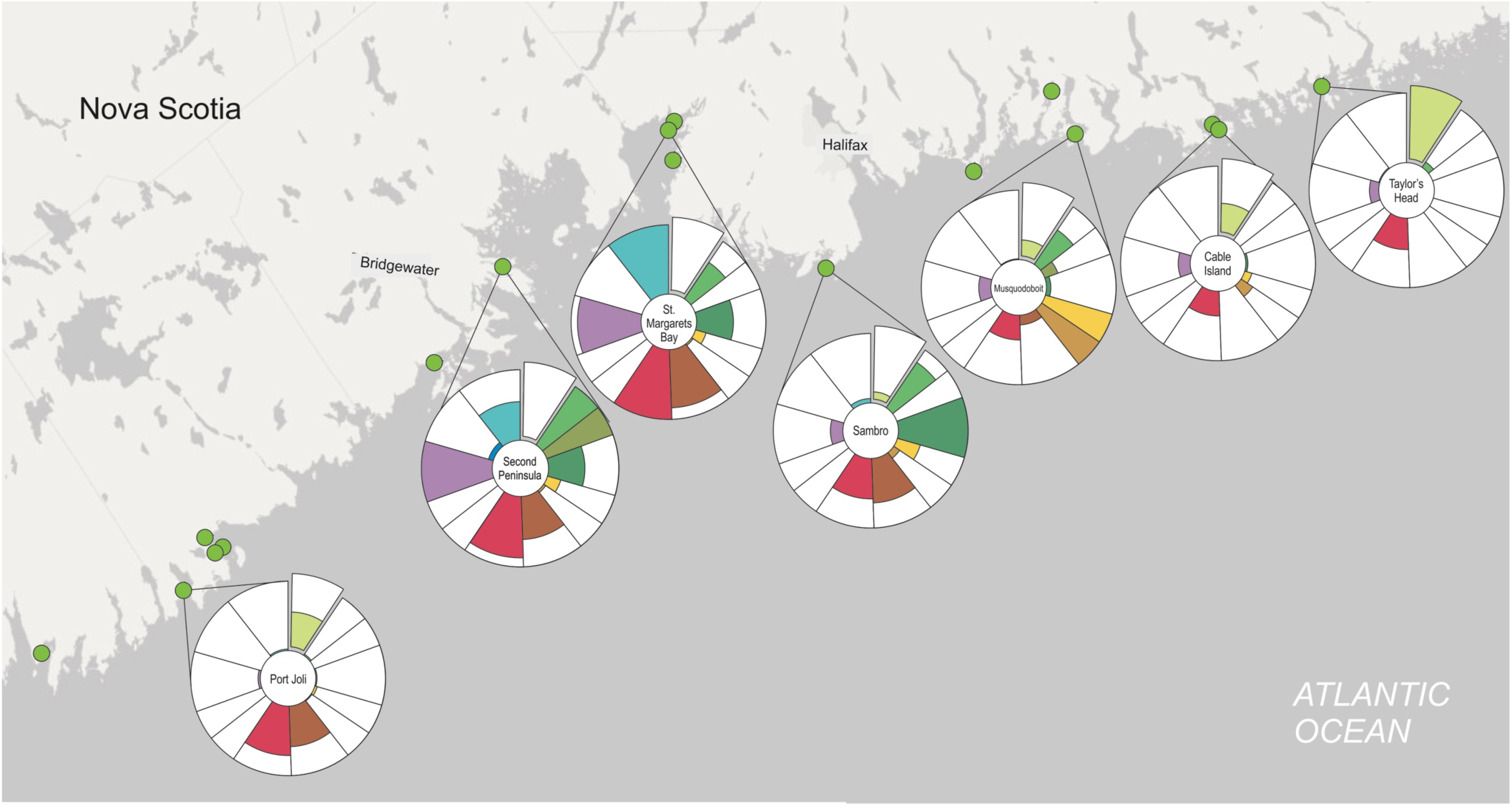
Standardized human impact metric for seagrass beds on the Atlantic coast of Nova Scotia. Each coloured petal represents one human impact measure with the outer ring of each circle representing the maximum possible impact score (i.e. petals that fill up more space represent a higher score of that impact). Impact scores are shown for 7 selected sites relative to all 17 sites (green points) in Atlantic NS. The human impact metric was calculated for all Atlantic NS region sites but is only shown for 7 selected sites. Refer to Fig. 1 for legend.

**Figure 6.**
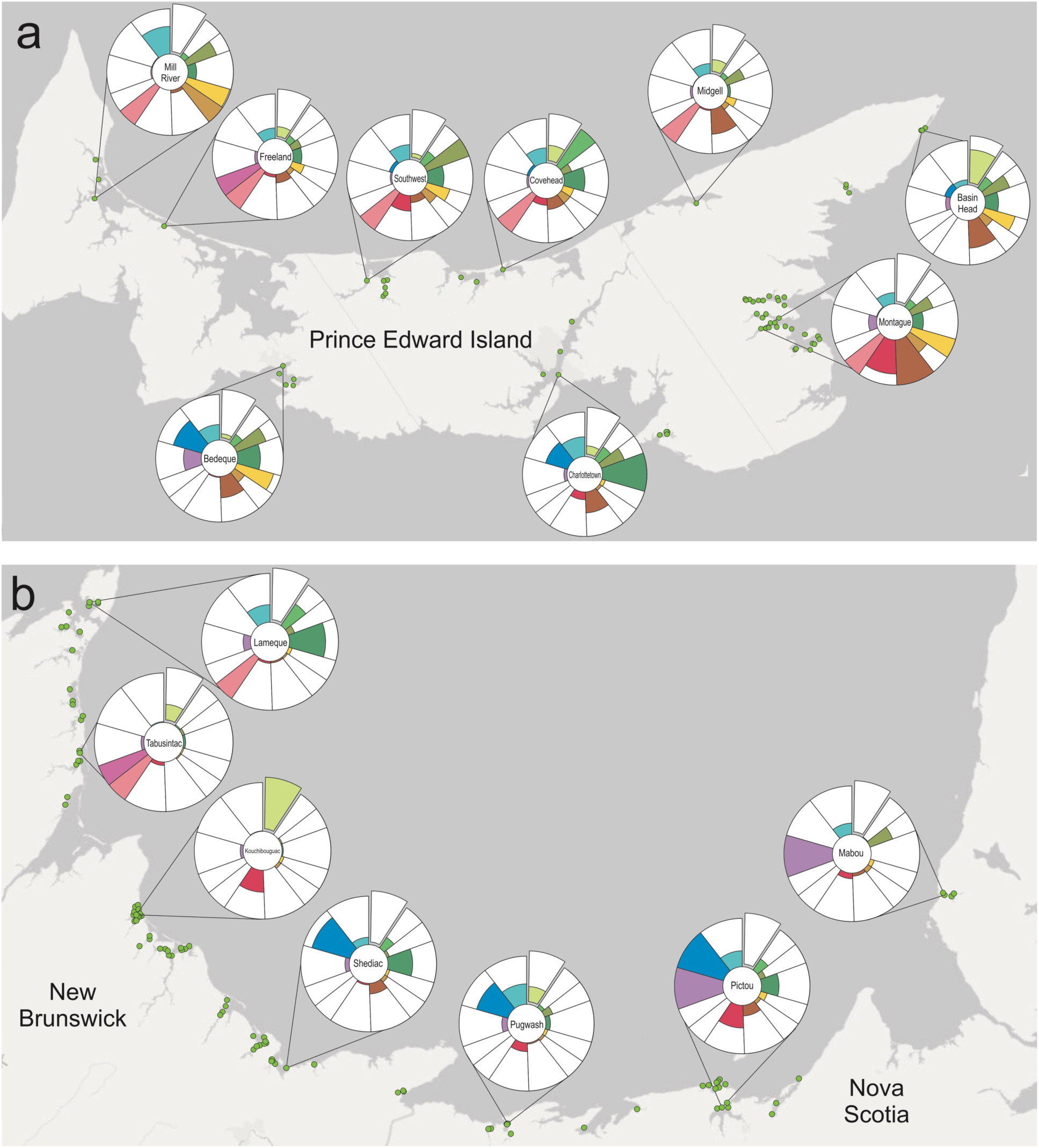
Standardized human impact metric for seagrass beds in (a) Prince Edward Island and (b) the Gulf coast of New Brunswick and Nova Scotia. Each coloured petal represents one human impact measure with the outer ring of each circle representing the maximum possible impact score (i.e. petals that fill up more space represent a higher score of that impact). Impact scores are shown for 9 sites in PEI and 7 in NB and NS but were calculated relative to all other 163 sites (green points) in the Gulf region. The human impact metric was calculated for all Gulf region sites but is only shown for 15 selected sites. Refer to Fig. 1 for legend.

Our human impact metric can also be used to assess the degree of impacts among individual seagrass beds within a bay. For example, Pugwash Bay (Fig. 7a) had relatively low human impacts at the bay-scale (i.e., almost all impacts within the 20-40^th^ percentile), but high heterogeneity in local-scale impacts among the individual seagrass beds. For example, Bed “C” (Fig. 7a) was only minimally impacted by poor water quality and overwater structures, while beds “A” and “B” were more highly impacted by overwater structures and reduced buffer zones. In comparison, Bedeque Bay in PEI (Fig. 7b) was highly impacted by human activities at the bay-scale (i.e., most bay-scale impacts in the 80-98^th^ percentile) with high agricultural land use and nitrogen loading. Again, high heterogeneity in human impacts was evident among the individual seagrass beds (Fig. 7b). For example, Bed “A” had extensive overwater structures in its vicinity and poor water quality, while beds ‘B’ and ‘C’ were influenced by a relatively low degree of human impacts at the local-scale.

**Figure 7.**
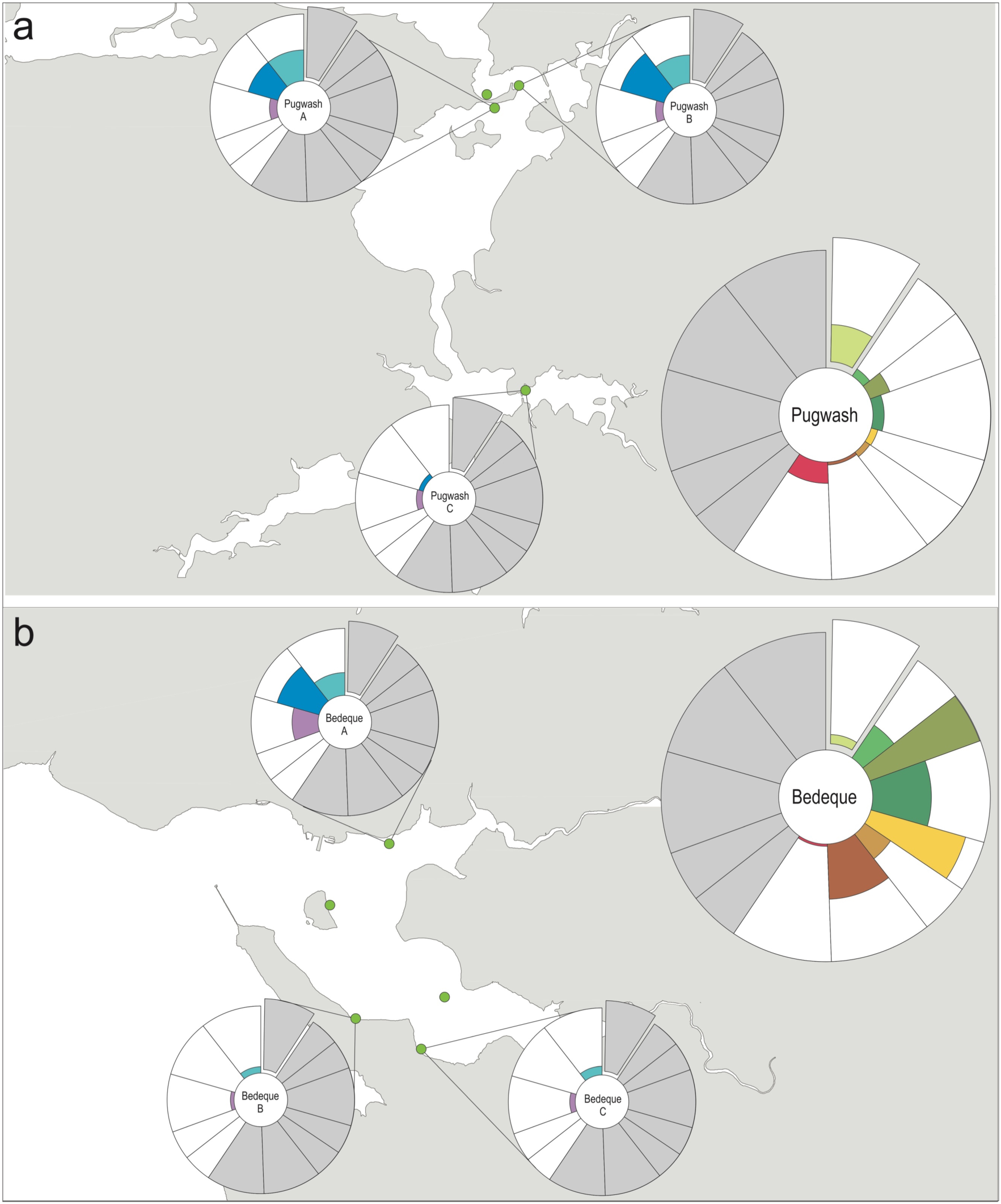
Standardized human impact metric for seagrass beds in (a) Pugwash Bay, NS and (b) Bedeque Bay, PEI in the Gulf region. The large petal diagrams in the corner of each plot represent the bay-scale impacts and the small petal diagrams represent the local-scale impacts for each individual bed, with the respective other impacts removed (grey petals). Each coloured petal represents one human impact measure with the outer ring of each circle representing the maximum possible impact score (i.e. petals that fill up more space represent a higher score of that impact). Impact scores are relative to all other sites in the Gulf region of NB, NS, and PEI. Green points indicate the locations of seagrass beds. Refer to Fig. 1 for legend.

## Discussion

Quantifying the magnitude and range of human impacts in a standardized, comprehensive and comparable way is urgently needed for large-scale scientific assessments and for management and conservation planning in coastal ecosystems. Previous metrics of human impacts for marine ecosystems have typically been developed on global, national, or broad regional scales, with little effort towards metrics appropriate for coastal ecosystems or specific biogenic habitats. Our human impact metric incorporates a suite of anthropogenic stressors from both terrestrial and marine sources relevant for coastal bays and estuaries as well as biogenic habitats and can be adapted to and standardized across multiple spatial scales.

Despite including multiple human impacts into our metric, we did not to combine them into one cumulative impact score (*sensu* Halpern *et al.* 2008), as this requires scientifically-informed vulnerability weightings that are not readily available. Although research on human impacts to seagrass ecosystems in Atlantic Canada has increased over the past decade (Coll *et al.* 2011, Schmidt *et al.* 2012, Hitchcock *et al.* 2017, Cullain *et al.* 2018a, McIver *et al.* in review), we still know too little about the responses of seagrass ecosystems to individual and cumulative human impacts to allow effective ranking or evidence-based weighting of their importance. Despite not explicitly accounting for the magnitude and relative importance of all impacts combined, our approach still provides a useful overall qualitative summary of the total human impact as visualized by the petal diagrams. Furthermore, providing individual impact scores instead of collapsing these into a single value allows influential impacts to be identified for specific bays, the relative impact to be compared, and also allows managers to identify and track impacts of interest. In the following, we discuss our application of the human impact metric to seagrass beds in Atlantic Canada, and then its potential for more general application to other habitats and regions and uses for management and conservation planning.

### Application to seagrass beds in Atlantic Canada

Applying our human impact metric to 180 seagrass beds in 52 coastal bays in Atlantic Canada revealed several insights relevant for scientific assessment, management and conservation. We found considerable regional variation in impact strength that provide insight into where coastal bays and seagrass ecosystems are expected to be most and least affected by individual or cumulative human impacts. Also, our results highlight the importance of considering human impacts at multiple spatial scales when assessing threats to coastal ecosystems and biogenic habitats.

One of the most damaging human impacts to the extent and condition of seagrass beds is eutrophication from human-derived nitrogen loading (Orth *et al.* 2006, Waycott *et al.* 2009). We found that 64% of all bays in our study are at risk of seagrass decline based on nitrogen loading rates (Latimer and Rego 2010). However, only 13% of bays, most of which are in PEI, are at risk for anoxic events based on ΔN values (Bugden *et al.* 2014). While both nitrogen loading and ΔN can indicate eutrophication risk, ΔN can indicate the potential for more severe anoxic events to occur by accounting for dilution from freshwater input and tidal flushing (Bugden *et al.* 2014). Thus, while human-derived nitrogen loading can be high in many bays and lead to primary (e.g. enhanced phytoplankton, epiphytic or benthic algal growth) or secondary (e.g. reduced shoot density and biomass) eutrophication effects in seagrass beds (Schmidt *et al.* 2012, Cullain *et al.* 2018a, McIver *et al.* in review), more severe loss of seagrass cover from anoxic events may only be of concern in PEI (Bugden *et al*. 2014).

Despite relatively low land use change within Atlantic NS watersheds, the wide range of riparian land alteration in close proximity to seagrass beds suggests that local-scale impacts may be important for seagrass health in Atlantic NS. Additionally, 42% of Atlantic NS bays have no coastal land protection. This is similar to bays in Gulf NB+NS but is much higher than PEI where only 11% of bays have no coastal land protection. Seagrass beds bordered by land managed for conservation purposes usually have higher temporal stability than beds associated with unprotected land (Breininger *et al*. 2016). Not surprisingly, our results show that coastal land protection at the bay-scale correlates well with the extent of riparian land alteration adjacent to seagrass beds i.e. local-scale (p < 0.001, R^2^ = 0.42, Fig. S3). This suggests that implementation of coastal conservation areas at the bay-scale may be a useful strategy to reduce the number of seagrass beds at risk of reduced buffer zones from riparian land alteration (Quiros *et al.* 2017).

Compared to the Gulf region, Atlantic NS bays have a higher extent of invasive biofouling species. This is not surprising given that the closer proximity of Atlantic NS to the open Atlantic Ocean and United States allows greater susceptibility to range expansions of invaders (Sephton *et al.* 2017). The difference in invasion extent between the two regions is driven by recent invasion of several tunicate species in NS that are absent from the Gulf (Moore *et al.* 2014, Vercaemer *et al.* 2015). However, species distribution models predict northward range expansion of most invasive tunicates, with many seagrass inhabited bays in the Gulf region becoming hotspots for invasive biofouling species over the next 50 years (Lowen and DiBacco 2017). Fouling by invasive tunicates reduces seagrass growth and survival due to shading and premature breakage (Brush and Nixon 2002, Wong and Vercaemer 2012). Since our human impact metric identified many highly impacted seagrass beds in the Gulf region, investigating how seagrass beds respond to the cumulative effect of invasive biofouling species and other human impacts should be a research priority since this may become a reality in the near future.

Overwater structures are recognized as having one of the largest negative impacts on seagrasses, yet no threshold level has been determined (Fresh et al. 2001, Rehr et al. 2014). Therefore, it remains unclear what degree of overwater structure coverage is detrimental to seagrass and if the range of impact scores we observed in Atlantic Canada are considered harmful. While the median percentage of water covered by overwater structures was relatively low for most seagrass beds, with no structures within 1km of 47% of seagrass beds, our impact metric identified several beds where coverage by overwater structures was much higher. These were primarily in the Gulf region and, in some cases, in bays where other human impacts were low. For example, overwater structures near seagrass beds in Pugwash Bay and Shediac Bay are relatively high despite low impact scores for most other threats (Fig. 6b). This illustrates the importance of quantifying numerous threats at multiple scales to fully understand impacts to biogenic habitats. Future research that identifies thresholds for the effects of overwater structures on coastal biogenic habitats would improve assessments of ecosystem health.

### Assessment of human impact at multiple spatial scales

Our human impact metric is unique as it assesses impacts at multiple spatial scales. This allows evaluation of within-bay variation and can reveal discrepancies about the relative magnitude of impacts at different spatial scales. A metric that accounts for different spatial scales of impact is critical for biogenic habitats in coastal ecosystems given the uncertainty in what human impacts, and at what spatial scales these impacts, are most detrimental to these habitats. Seagrass responses to anthropogenic stressors typically exhibit a high level of within-bay variability (Hitchcock *et al.* 2017), and our results show that there is also often high within-bay variation in stressor intensity (Fig. 7). By considering impacts at both local and bay-wide scales our metric can highlight locations where there is spatial mismatch between the intensity of bay-wide and local-scale impacts at individual seagrass beds. For example, Bedeque Bay, PEI is a highly impacted bay based on bay-scale impacts, however, there is high variability among local-scale impacts within the bay (Fig. 7b). Observation of only local-scale impacts would suggest certain beds are relatively non-impacted when this might not be the case. This emphasizes the necessity of quantifying impacts at multiple scales to fully understand the degree of impact to coastal biogenic habitats.

### General application to other habitats and ecosystems

Along with seagrass, several other species create complex biogenic habitats in coastal waters in Canada and elsewhere, including rockweed beds, kelp forests, oyster reefs, and others (Beazley *et al.* 2017). Despite broad ideas of how these biogenic habitats respond to human pressures, little is known of the trajectories for individual habitats, what management actions are needed, and where to prioritize conservation efforts (DFO 2018). When developing our human impact metric, we focused on impacts that are not only relevant for biogenic habitats but where similar data to quantify the extent of each impact is obtainable for other provinces and coasts in Canada, as well as other countries. For example, the data used to calculate water quality, coastal fishing activity, and invasion extent are all from federal programs that collect comparable data for other coastal regions of Canada. The model used to estimate nitrogen loading rates can be adapted to coastal bays and estuaries worldwide (Valiela *et al.* 1997), and the data used to calculate coastal land protection, land uses, human population density, and aquaculture presence are all provincially regulated with comparable data available across provinces. Thus, our human impact metric can be readily applied to other coastal regions and habitats in Canada and beyond. This opens the door for direct comparisons of human impacts to a specific habitat, i.e. seagrass beds, across provinces, biogeographic regions, and coasts, as well as comparisons between different biogenic habitats within and across countries.

The raw and standardized impact scores for all 180 seagrass beds and 52 bays assessed in this study (data appendix A and B) are available to aid in the development of monitoring programs, management strategies and prioritization of coastal conservation efforts in Atlantic Canada. This database will be valuable for the application of our human impact metric to other biogenic coastal ecosystems in Atlantic Canada, as many of the seagrass inhabited bays are also home to kelp and rockweed habitats. Therefore, the same bay-scale impacts already calculated for the current database apply and only local-scale impacts need to be adjusted.

Our impact metric has been specifically tailored for impacts relevant to seagrass beds in Atlantic Canada. Applications to other ecosystems and regions should begin with an assessment of the impacts present and their potential influence (e.g. Table 1). For example, boat moorings are not an important threat to Atlantic Canada seagrass beds yet should be included in impact metrics for seagrass in parts of the coastal United States, United Kingdom, and Australia (Hallac *et al.* 2012, Demers *et al.* 2013, Unsworth *et al.* 2017). Also, some impacts may be more relevant for one habitat type than another within the same region, such as commercial plant harvesting in Atlantic Canada rockweed beds (DFO 2013). Furthermore, our human impact metric does not consider climate change related stressors (i.e. increased sea surface temperature and sea level rise), which impact seagrass through loss of suitable habitat and direct effects on survival (Waycott *et al.* 2007, Olsen *et al.* 2012, Valle *et al.* 2014). Large-scale climate change impacts were not relevant for the application of our metric to seagrass beds in Atlantic Canada but could be included for more spatially broad applications.

### Application to management and conservation

Our metric is valuable for conservation planning by highlighting areas of low impact and high naturalness, which is a conservation priority for marine protected area placement in Canada and around the world (DFO 2004, Yamakita *et al.* 2015). For example, in the Scotian Shelf our impact metric highlights three seagrass beds in the Eastern Shore Archipelago with a high degree of coastal land protection and among the lowest impact scores relative to other beds in this region (Fig. 5, Table S1 and S2). This provides evidence for the Eastern Shore Archipelago as an ideal candidate for marine protection given the low human impact and high naturalness at both the bay scale and in close proximity to a valuable biogenic habitat (DFO in press). By assessing impacts in both the terrestrial and marine realms our impact metric will also be valuable for land-sea conservation planning by identifying coastal ecosystems adjacent to protected land (Alvarez-Romero *et al.* 2015). For example, in the Scotian Shelf, Port Joli has high coastal land protection and low riparian land alteration relative to other bays in the region (Fig. 5, Table S1 and S2). In the Gulf, Kouchibouguac and St. Louis de Kent (Fig. 6, Table S3) have 100% coastal land protection due to the presence of a National Park.

The comprehensive, standardized, and multi-scale nature of our metric can also assist in developing management strategies for individual or multiple human impacts. For example, in the Scotian Shelf, Second Peninsula ranks among the most highly impacted sites (Fig. 5, Table S1 and S2) with nitrogen loading close to the threshold detrimental to seagrass cover (Latimer and Rego 2010), high levels of land use that exceed the threshold for risk of ecosystem degradation in receiving waters (WWF 2017), poor water quality, and no coastal land conservation. Thus, our impact metric identifies risks from multiple human impacts that could be addressed by specific management strategies.

The impacts can also be linked to key mechanisms of growth and survival. For example, light availability is a key determinant of seagrass condition (Hauxwell *et al.* 2003), thus, knowledge of which seagrass beds are subject to multiple light limiting impacts will be beneficial for management. In the Gulf, our metric identified several seagrass beds at risk from multiple forms of light limitation. For example, nitrogen loading for Bedeque PEI is among the highest in the Gulf region (Table S3), and one seagrass bed in Bedeque Bay also has among the highest impact scores for overwater structures and riparian land alteration (Table S4). The combination of bay- and local-scale impacts specifically influencing light availability suggests that this bed may be an ideal candidate for management actions targeted at maintaining water clarity, such as reductions in land-derived nutrient inputs (Leschen *et al.* 2010, Greening *et al.* 2014, Lefchek *et al.* 2018), removal of overwater structures (Thom *et al.* 2005, Rehr *et al.* 2014), and conservation of coastal land (Stoms *et al.* 2005, Alvarez-Romero *et al.* 2015).

### Caveats and the way forward

One current limitation of our metric is that temporal variation could only be accounted for with some impacts (nutrient pollution, invasion extent, and water quality), which represent 10-year averages. The remaining impacts reflect intensity at the present time. This may be problematic if impacts been historically higher but are currently reduced through management strategies. Inclusion of temporal impact data would be useful to evaluate the response time of biogenic habitats to impacts and should be included where possible.

Our application of the human impact metric is also limited by the disparity in spatial distribution of seagrass sites among regions (n = 163 and 17 for Gulf and Scotian Shelf regions, respectively). Seagrass beds included in our study were those sampled in summer field surveys over the past decade. Inclusion of sites from a large-scale monitoring program in the Gulf region (e.g. Community Aquatic Monitoring Program; Weldon et al. 2009) resulted in a much larger sample size relative to Atlantic NS, where data were compiled from small, individual research projects. The larger dataset in the Gulf region highlighted the importance of within-bay variability, which could not be assessed for the Scotian Shelf region. Further application of our human impact metric as more studies are conducted would be beneficial to fully map the distribution of anthropogenic stressors in this region.

Finally, further work is necessary to allow the individual and combined impacts to be ranked according to importance for seagrass health. An understanding of the relative importance of each impact will allow the calculation of cumulative scores based on vulnerability weightings, producing one overall metric that is easily interpretable and useful for management and conservation planning.

## Conclusions

Given that coastal ecosystems are highly susceptible to human activities, the framework we have developed for quantifying human impacts is valuable for conservation and management planning. Our impact metric allows the finer-scale dynamics of anthropogenic stressors in coastal ecosystems to be explored by uniquely focusing on stressors relevant for coastal biogenic habitats across two spatial scales. Our metric can be applied to coastal ecosystems worldwide and can be used to prioritize areas for protection and management by identifying areas of low and high impact, highlighting prominent impacts within and between regions, and describing within-bay variation. The application of our human impact metric to seagrass beds across Atlantic Canada reveals a gradient of human impacts and identifies several bays and seagrass beds that may be good candidates for protection based on high naturalness, and others at risk for future degradation if management strategies are not implemented. Our results can also be used to apply the impact metric to other biogenic habitats in Atlantic Canada and as a baseline of human activities for future comparisons.

## Acknowledgments

This research is sponsored by the NSERC Canadian Healthy Oceans Network and its Partners: Department of Fisheries and Oceans Canada and INREST (representing the Port of Sept-Îles and City of Sept-Îles). We would like to thank Marc Skinner, Renee Bernier, Andrea Locke, Monica Boudreau and the DFO Community Aquatic Monitoring Program for seagrass location data. Thanks to the Canadian Shellfish Sanitation Program for supplying water quality data, the DFO Aquatic Invasive Species Program for supplying invasive species data, Adam Cook for supplying lobster fishing data, and Elizabeth Nagel for assistance with data collection.

## References

Alvarez-Romero JG, Pressey RL, Ban NC, and Brodie J. 2015. Advancing land-sea conservation planning: Integrated modelling of catchments, land-use change, and river plumes to prioritise catchment management and protections. PLoS ONE, 10: e0145574.

Anderson SC, Mills Flemming J, Watson R, and Lotze HK. 2011. Rapid global expansion of invertebrate fisheries: Trends, Drivers, and Ecoystem Effects. PLoS ONE, 6:e14735.

Arasamuthu A, Mathews G, and Patterson Edward JK. 2017. Spatial differences in bacterial and water quality parameters in seagrass meadows of Tutucorin Coast, Gulf of Mannar, southeastern India. Journal of Aquatic Biology and Fisheries, 5:1–10.

Ban N and Alder J. 2008. How wild is the ocean? Assessing the intensity of anthropogenic marine activities in British Columbia, Canada. Aquatic Conservation: Marine and Freshwater Ecosystems, 18:55–85.

Bastien-Daigle S, Hardy M, and Robichaud G. 2007. Habitat management qualitative risk assessment: Water column oyster aquaculture in New Brunswick. Canadian Technical Report of Fisheries and Aquatic Sciences, 2728: vii + 72p.

Beazley L, Kenchington E, and Lirette C. 2017. Species distribution modelling and kernel density analysis of benthic ecologically and biologically significant areas (EBSAs) and other benthic fauna in the Maritimes Region. Canadian Technical Report of Fisheries and Aquatic Sciences, 3204.

Beck MW, Brumbaugh RD, Airoldi L, Carranza A, Coen LD, Crawford C, et al. 2011. Oyster reefs at risk and recommendations for conservation, restoration, and management. Bioscience, 61:107–116.

Bilkovic DM, Roggero M, Hershner CH, and Havens KH. 2006. Influence of land use on microbenthic communities in nearshore estuarine habitats. Estuaries and Coasts, 29:1185–1195.

Blake RE, Duffy JE, and Richardson JP. 2014. Patterns of seagrass community response to local shoreline development. Estuaries and Coasts, 37:1549–1561.

Breininger DR, Breininger RD, and Hall CR. 2016. Effects of surrounding land use and water depth on seagrass dynamics relative to a catastrophic algal bloom. Conservation Biology 31:6775.

Bryce SA, Lomnicky GA, and Kaufmann PR. 2010. Protecting sediment-sensitive aquatic species in mountain streams through the application of biologically based sediment criteria. Journal of the North American Benthological Society, 29:657–672.

Bugden G, Jiang Y, van den Heuvel M, Vandermeulen H, MacQuarrie K, Crane C, et al. 2014. Nitrogen loading criteria for estuaries in Prince Edward Island. Canadian Technical Report of Fisheries and Aquatic Sciences, 3066:3(1).

Braga RR, Gomez-Aparicio L, Heger T, Vitule JRS, Jeschke JM. 2018. Structuring evidence for invasional meltdown: broad support but with biases and gaps. Biological Invasions, 20:923–936.

Brush ML and Nixon SW. 2002. Direct measurements of light attenuation by epiphytes on eelgrass *Zostera marina*. Marine Ecology Progress Series, 238: 73–79.

Burgin S and Hardiman N. 2011. The direct physical, chemical, and biotic impacts on Australian coatal waters due to recreational boating. Biodiversity and Conservation, 20:683–701.

Coll M, Schmidt A, Romanuk T and Lotze HK. 2011. Food-web structure of seagrass communities across different spatial scales and human impacts. PLoS One, 6:e22591

Comeau, LA. 2013. Suspended versus bottom oyster culture in eastern Canada: Coparing stocking densities and clearance rates. Aquaculture, 410, 57–65.

CSSP. 2016. Available from: http://www.inspection.gc.ca/food/fish-and-seafood/shellfish-sanitation/eng/1299826806807/1299826912745

Cullen-Unsworth LC and Unsworth RKF. 2016. Strategies to enhance the resilience of the world’s seagrass meadows. Journal of Applied Ecology, 53:967–972

Cullain N, McIver R, Schmidt AL, and Lotze HK. 2018a. Impacts of organic enrichment from finfish aquaculture on seagrass beds and associated macroinfaunal communities in Atlantic Canada. PeerJ (in press)

Cullain N, McIver R, Schmidt AL, and Lotze HK. 2018b. Spatial variation of macroinfaunal communities associated with *Zostera marina* beds across three biogeographic regions in Atlantic Canada. Estuaries and Coasts, 41:1381–1396.

Demers M-CA, Davis AR, and Knott NA. 2013. A comparison of the impact of ‘seagrass-friendly’ boat mooring systems on *Posidonia australis*. Marine Environmental Research, 83:5462.

Department of Fisheries and Oceans Canada. 2004. Evaluation of site selection methodologies for use in marine protected area network design. Canadian Science Advisory Secretariat Science Advisory Report, 2004/082.

Department of Fisheries and Oceans Canada. 2007. Ecologically and Biologically Significant Areas (EBSA) in the Estuary and Gulf of St. Lawrence: identification and characterization. Canadian Science Advisory Secretariat Science Advisory Report, 2007/016.

Department of Fisheries and Oceans Canada. 2009a. Does eelgrass (*Zostera marina*) meet the criteria as an ecologically significant species? CSAS Science Advisory Report, 2009/018.

Department of Fisheries and Oceans Canada. 2009b. Development of a framework and principles for the biogeographic classification of Canadian marine areas. CSAS Science Advisory Report, 2009/056.

Department of Fisheries and Oceans Canada. 2013. Assessment of information on Irish moss, rockweed, and kelp harvests in Nova Scotia. CSAS Science Advisory Report, 2013/004.

Department of Fisheries and Oceans Canada. 2018. Design strategies for a network of marine protected areas in the scotia shelf bioregion. CSAS Science Advisory Report, 2018/006

Department of Fisheries and Oceans Canada. In press. Biophysical and Ecological Overview of the Eastern Shore Islands Area of Interest (AOI). CSAS Science Advisory Report, 2018/

Duarte CM. 2002. The future of seagrass meadows. Environmental Conservation, 29:192–206.

Google Inc. 2018. Google Earth Pro (Version 7.3.2.5491) [Software].

Fresh KL, Wyllie-Echeverria T, Wyllie-Echeverria S, and Williams BW. 2006. Using light-permeable grating to mitigate impacts of residential floats on eelgrass *Zostera marina* L. in Puget Sound, Washington. Ecological Engineering, 28:354–362.

Garbary DJ, Miller AG, Williams, J, Seymour, NR. 2014. Drastic decline of an extensive eelgrass bed in Nova Scotia due to the activity of the invasive green crab (Carcinus maenas). Marine Biology, 161:3–15.

Grech A, Chartrand-Miller K, Erftemeijer P, Fonseca M, McKenzie L, Rasheed M, et al. 2012. A comparison of threats, vulnerabilities and management approaches in global seagrass bioregions. Environmental Research Letters, 7:024006.

Greening H, Janicki A, Sherwood ET, Pribble R, and Johansson JOR. 2014. Ecosystem responses to long-term nutrient management in an urban estuary: Tampa Bay, Florida, USA. Estuarine, Coastal, and Shelf Science, 151:1–16.

Guyondet T, Sonier R, and Comeau LA. 2013. A spatially explicit seston depletion index to optimize shellfish culture. Aquaculture Environment Interactions, 4:175–186.

Hallac D, Sadle J, Pearlstine LG, Herling F, and Shinde D. 2012. Boating impacts to seagrass in Florida Bay, Everglades National Park, Florida, USA: links with physical and visitor-use factors and implications for managaement. Marine and Freshwater Research, 6:1117–1128.

Halpern BS, Walbridge S, Selkoe KA, Kappel CV, Micheli F, D’agrosa, et al. 2008. A global map of human impact on marine ecosystems. Science, 319:948–953.

Hauxwell J, Cebrián J, and Valiela I. 2003. Eelgrass *Zostera marina* loss in temperate estuaries: Relationship to land-derived nitrogen loads and effect of light limitation imposed by algae. Marine Ecology Progress Series, 247:59–73.

Hemminga MA, and Duarte CM. 2000. Seagrass Ecology. Cambridge University Press.

Hitchcock JK, Courtenay SC, Coffin MRS, Pater CC, and van ven Heuval MR. 2017. Eelgrass bed structure, leaf nutrient, and leaf isotope responses to natural and anthropogenic gradients in estuaries of the southern Gulf of St. Lawrence, Canada. Estuaries and Coasts, 40:1653–1665.

Huang J, Huang Y, Pontius RG, and Zhang Z. 2015. Geographically weighted regression to measure spatial variations in correlations between water pollution versus land use in a coastal watershed. Ocean and Coastal Management 103:14–24.

Hughes AR, Williams SL, Duarte CM, Heck KL, Waycott M, 2009. Associations of concern: declining seagrasses and threatened dependent species. Frontiers in Ecology and the Environment, 7:242–246.

Iacerella JC, Adamczyk E, Bowen D, Chalifour L, Eger A, Heath W, et al. 2018. Anthropogenic disturbance homogenizes seagrass fish communities. Global Change Biology, 00:1–15.

Krumhansl KA, Okamoto DK, Rassweiler A, Novak M, Bolton JJ, Cavanaugh KC, et al. 2016. Global patterns of kelp forest change over the past half-century. Proceedings of the National Academy of Sciences, 113:13785–13790.

Lamb JB, van de Water JAJM, Bourne DG, Altier C, Hein MY, Fiorenza EA, et al. 2017. Seagrass ecosystems reduce exposure to bacterial pathogens of humans, fishes, and invertebrates. Science, 355:731–733.

Latimer JS and Rego SA. 2010. Empirical relationship between eelgrass extent and predicted watershed-derived nitrogen loading for shallow New England estuaries. Estuarine, Coastal and Shelf Science, 90:231–240.

Lefcheck JS, Orth RJ, Dennison WC, Wilcox DJ, Murphy RR, Keisman J, et al. 2018. Long-term nutrient reductions lead to the unprecedented recovery of a temperate coastal region. Proceedings of the National Academy of Sciences, 115:3658–3662.

Leitao RP, Zuanon J, Mouillot D, Leal CG, Hughes RM, Kaufmann PR, et al. 2018. Disentangling the pathways of land use impacts on the functional structure of fish assemblages in Amazon streams. Ecography, 41:219–232.

Lerberg SB, Holland AF, and Sanger DM. 2000. Responses of tidal creek microbenthic communities to the effects of watershed development. Estuaries, 23:838–853.

Leschen AS, Ford KH, and Evans T. 2010. Successful eelgrass (Zostera marina) restoration ina. Formerly eutrophic estuary (Boston Harbor) supports the use of a multifaceted watershed approach to mitigating eelgrass loss. Estuaries and Coasts, 33:1340–1354.

Lotze HK, Lenihan HS, Bourque BJ, Bradbury RH, Cooke RG, Kay MC, et al. 2006. Depletion, degradation, and recovery potential of estuaries and coastal seas. Science, 312:1806–1809.

Lowen JB and DiBacco C. 2017. Distributional changes in a guild of non-indigenous tunicates in the NW Atlantic under high-resolution climate projections. Marine Ecology Progress Series, 570:173–186.

McIver R, Milewski I, and Lotze HK. 2015. Land use and nitrogen loading in seven estuaries along the southern Gulf of St. Lawrence, Canada. Estuarine, Coastal and Shelf Science, 165:137–148.

McIver R, Milewski I, Loucks R, and Smith R. 2018. Estimating nitrogen loading and far-field dispersal potential from background sources and coastal finfish aquaculture: A simple framework and case study in Atlantic Canada. Estuarine, Coastal and Shelf Science, 205:46–57.

McIver R, Schmidt AL, Cullain N, and Lotze HK. In review. Linking estimates of nitrogen loading and watershed characteristics to eelgrass bed structure and eutrophication symptoms across 7 bays in Atlantic Canada. Marine Environmental Research (in revision MERE_2017_577).

Moore A, Vercaemer B, DiBacco C, Sephton D, and Ma K. 2014. Invading Nova Scotia: first records of *Didemnum vexillum* Kott, 2002 and four more non-indigenous invertebrates in 2012 and 2013. BioInvasions Records, 3:225–234.

Murray CC, Agbayani S, Alidini HM, and Ban NC. 2015. Advancing marine cumulative effects mapping: An update in Canada’s Pacific waters. Marine Policy, 58:71–77.

Nagel EJ, Murphy G, Wong MC and Lotze HK. 2018. Nitrogen loading rates for twenty-one seagrass inhabited bays in Nova Scotia, Canada. Canadian Technical Report of Fisheries and Aquatic Sciences, 3260: v + 37.

Olsen YS, Sanchez-Camacho M, Marba N, and Duarte CM. 2012. Mediterranean seagrass growth and demography responses to experimental warming. Estuaries and Coasts, 35:12051213.

Orth RJ, Carruthers TJB, Dennison WC, Duarte CM, Fourqurean JW, Heck KLJr, et al. 2006. A global crisis for seagrass ecosystems. Bioscience, 56:987–996.

Palmer T. 2018. Nitrogen loading rates for Prince Edward Island estuaries and effects on eelgrass (Zostera marina) structural characteristics. Honours Thesis, Dalhousie University.

Parker J, Epifanio J, Casper A, and Cao Y. 2016. The effects of improved water quality on fish assemblages in a heavily modified large river system. River Research and Applications, 32:992–1007.

Powell KI, Chase JM, and Knight TM. 2013. Invasive plants have scale-dependent effects on diversity by altering species-area relationships. Science 339:316–318.

Quiros TEA, 2016. Linking Terrestrial and Marine Protected Areas at the Coastal Interface. PhD. University of Califiornia, Santa Cruz.

Quiros TEA, Croll D, Tershy C, Fortes MD, Raimondi P. 2017. Land use is a better predictor of tropical seagrass condition than marine protection. Biological Conservation 209:454–463.

Rehr AP, Williams GD, Tolimeieri N, and Levin PS. 2014. Impacts of terrestrial and shoreline stressors on eelgrass in Puget Sount: An expert elicitation. Coastal Management, 42:246–262.

Schmidt AL, Wysmyk JKC, Craig SE, and Lotze HK. 2012. Regional-scale effects of eutrophication on ecosystem structure and services of seagrass beds. Limnology and Oceanography, 57:1389–1402.

Sephton D, Vercaemer B, Silva A, Stiles, L, Harris M, and Godin K. 2017. Biofouling monitoring for aquatic invasive species (AIS) in DFO Maritimes Region (Atlantic shore of Nova Scotia and southwest New Brunswick): May - November, 2012 - 2015. Canadian Technical Report of Fisheries and Aquatic Sciences, 3158: ix + 172p.

Shelton AO, Francis TB, Feist BE, Williams GD, Lindquist A, and Levin PS. 2017. Forty years of seagrass population stability and resilience in an urbanizing estuary. Journal of Ecology, 105:458–470.

Short FT, and Wyllie-Echeverria, S. 1996. Natural and human-induced disturbance of seagrasses. Environmental Conservation, 23:17–27.

Simberloff D and Von Holle B. 1999. Positive interactions of nonindigenous species: Invasional meltdown? Biological Invasions, 1:21–32.

Simpson SD, Radford AN, Nedelec SL, Ferrari MCO, Chivers DP, Mccormick MI, et al. 2016. Anthropogenic noise increases fish mortality by predation. Nature Communications, 7:10544.

Skinner MA, Courtenay SC, and McKindsey CW. 2013. Reductions in distribution, photosynthesis, and productivity of eelgrass Zostera marina associated with oyster Crassostrea virginica aquaculture. Marine Ecology Progress Series, 486:105–119.

Spalding MD, Fox HE, Allen GR, Davidson N, Ferdana ZA, Finlayson M, et al. 2007. Marine ecoregions of the world: a bioregionalization of coastal and shelf areas. Bioscience, 57:573–583.

Stoms DM, Davis FW, Andelman SJ, Carr MH, Gaines SD, Halpern BS, et al. 2005. Integrated coastal reserve planning: Making the land-sea connection. Frontiers in Ecology and the Environment, 3:429–436.

Thom RM, Williams GW, and Diefenderfer HL. 2005. Balancing the need to develop coastal areas with the desire for an ecologically functioning coastal environment: Is net ecosystem improvement possible. Restoration Ecology, 13:193–203.

Thom RM, Buenau KE, Judd C, and Cullinan VI. 2011. Eelgrass (Zostera marina) stressors in Puget Sound. U.S. Department of Energy.

Thrush SF, Lawrie SM, Hewitt JE, and Cummings VJ. 1999. The problem of scale: Uncertainties and implications for soft-bottom marine communities and the assessment of human impacts. Biogeochemical Cycling and Sediment Ecology, 59:195–210.

Uhrin AV, Fonesca MS, and DiDomenico GP. 2005. Effect of Caribbean spiny lobster traps on seagrass beds of the Florida Keys National Marine Sanctuary: Damage assessment and evalutation of recovery. American Fisheries Society Symposium 41:579–588.

Unsworth RKF, Williams B, Jones BL, and Cullen-Unsworth LC. Rocking the boat: Damage to eelgrass by swinging boat moorings. Frontiers in Plant Science. 8:1309.

Uriarte M, Yackulic CB, Lim Y, and Arce-Nazario JA. 2011. Influence of land use on water quality in a tropical landscape: a multi-scale analysis. Landscape Ecology, 26:1151–1164.

Valiela I, Collins G, Kremer J, Lajtha K, Geist M, Seely B, et al. 1997. Nitrogen loading from coastal watersheds to receiving estuaries: New method and application. Ecological Applications, 7:358–380.

Valle M, Chust G, del Campo A, Wisz MS, Olsen SM, Garmendia JM, et al. 2014. Projecting future distribution of the seagrass *Zostera noltii* under global warming and sea level rise. Biological Conservation, 170:74–85

Van Katwijk MM, van der Welle MEW, Lucassen ECHET, Vonk JA, Christianen MJA, et al. 2011. Early warning indicators for river nutrient and sediment loads in tropical seagrass beds: a benchmark from a near-pristine archipelago in Indonesia. Marine Pollution Bulletin, 62:1512-1520.

Vance A. 2014. Applying an ecosystem-based risk management approach to the relationship between eelgrass beds and oyster aquaculture at multiple spatial scales in eastern New Brunswick, Atlantic Canada. Master of Marine Management Thesis, University.

Vercaemer B, Sephton D, Clément P, Harman A, Stewart-Clark S, and Dibacco C. 2015. Distribution of the non-indigenous colonial ascidian *Didemnum vexillum* (Kott, 2002) in the Bay of Fundy and on offshore banks, eastern Canada. Managing Biological Invasions, 6:385–394.

Waycott M, Collier C, Mcmahon K, Ralph P, Mckenzie L, Udy J, et al. 2007. Vulnerability of seagrasses in the Great Barrier Reef to climate change. In J.E. Johnson, P.A. Marshal (Eds.), Climate Change and the Great Barrier Reef: A Vulnerability Assessment, Great Barrier Reef Marine Park Authority and Australian Greenhouse Office, Australia (2007), pp. 193–299.

Waycott M, Duarte CM, Carruthers TJB, Orth RJ, Dennison WC, Olyarnik S, et al. 2009. Accelerating loss of seagrasses across the globe threatens coastal ecosystems. Proceedings of the National Academy of Science, 106:12377–12381.

Weldon J, Courtenay S, and Garbary D. 2009. The community aquatic monitoring program (CAMP) for measuring marine environmental health in coastal waters of the southern Gulf of St. Lawrence: 2007 overview. Canadian Technical Report of Fisheries and Aquatic Sciences 2825.

Williams SL. 2007. Introduced species in seagrass ecosystems: Status and concerns. Journal of Experimental Marine Biology and Ecology, 350:89–110.

Wong MC. 2018. Secondary Production of Macrobenthic Communities in Seagrass (Zostera marina, Eelgrass) Beds and Bare Soft Sediments Across Differing Environmental Conditions in Atlantic Canada. Estuaries and Coasts, 41:536–548.

Wong MC and Vercaemer B. 2012. Effects of invasive colonial tunicates and a native sponge on the growth, survival, and light attenuation of eelgrass (*Zostera marina*). Aquatic Invasions, 7:315–326.

Worm B and Lotze HK. 2006. Eutrophication, grazing, and algal blooms on rocky shores. Limnology and Oceanography, 51:569–579.

World Wildlife Fund. 2017. A national assessment of Canada’s freshwater watershed reports. WWF Report.

Yamakita T, Yamamoto H, Nakaoka M, Yamano H, Fujukura K, Hidaka K, et al. 2015. Identification of importance marine areas around the Japanses Archipelago: Establishment of a protocol for evaluating a broad area using ecologically and biologically significant areas and selection criteria. Marine Policy, 51:136–147.

